# Apigenin exerts anti-cancer effects in colon cancer possibly by regulating Heat shock protein 90 alpha family class A member 1 (HSP90AA1)

**DOI:** 10.1101/2023.03.24.534119

**Authors:** Arindam Sain, Dipshikha Khamrai, Thirukumaran Kandasamy, Debdut Naskar

## Abstract

Apigenin, a natural flavonoid, has shown early promise in colon cancer (CC); thus, exploring potential mechanisms of apigenin in CC is obligatory. In this study, shared targets of apigenin and CC were identified through different online tools and subjected to functional enrichment analyses like Gene Ontology and KEGG. Further, the protein-protein interaction network of the shared targets was developed (via STRING); hub/core targets were identified (MCODE application). The top targets of apigenin in CC were identified by molecular docking; further investigated for differential gene and protein expression in CC and their influence on CC patient survival (using TCGA data). Based on the docking score of the 13 hub genes, the top 3 targets (*HSP90AA1, MMP9, PTGS2*) were selected, and their expression was significantly elevated and related to poor overall survival in CC (except *PTGS2*). Molecular dynamics simulation further validated protein-ligand interactions and selected HSP90AA1 as the best target of apigenin in CC. Finally, apigenin was found to be involved in the cytotoxicity of CC cells (COLO-205) by reducing *HSP90AA1* expression. The results of this study identified HSP90AA1 as one of the prime targets of apigenin in CC, and apigenin might act on HSP90AA1 to exert its anti-cancer mechanism.

## 1. INTRODUCTION

Colon cancer (CC) has become a leading threat in the healthcare system globally and significantly contributes to cancer mortality and morbidity around the globe [1]. Dietary habit (higher intake of red meat), sedentary lifestyle, genomic abnormalities, and augmented signalling pathways (linked to cellular growth and proliferation) dictate the progression of CC that mainly affect the entire large intestine or bowel [2]. Another alarming statistic shows that one in five colon cancer patients is aged 20-54 [3]. Moreover, in young adults, it develops as the third primary reason for cancer-related death [4].

Modifying the dietary pattern by consuming plant products such as phytochemicals has come a long way in the last decades and has shown promise in different cancers [5]. Changing lifestyle by doing proper physical activities, reducing extra fat, and continuing a healthy diet may prevent 40% of cancer cases [6]. On the other hand, the conventional way (chemotherapy and surgical resection) of treating or managing CC is still not optimum and adequate due to treatment-induced adverse effects [7]. Even rapidly developing therapeutic regimes, for example, immunotherapy, is beyond the reach of most patients due to high costs. Therefore, exploring alternative treatment/preventive strategies is obligatory to overcome the burden of costly treatment options and treatment-induced side effects.

Plant-based compounds like flavonoids, lignans, and tannins have been extensively studied for the last few decades in the search for novel cancer therapy [8]. Apigenin, a plant-derived edible flavonoid found in various plants (parsley, onions, celery, and chamomile), is one of the most studied phenols in cancer research [9]. Mainly due to its anti-cancer, antioxidant and anti-inflammatory activities in different cancer cells (e.g., breast, prostate, and colon cancer) [10]. The anti-cancerous properties of apigenin depend on its ability to modulate major growth signalling pathways (e.g., Akt/mTOR, NF-κB, STAT3) [11]. Additionally, apigenin diminished the Wnt/β-catenin pathway, significantly influencing CC pathogenesis [12].

Chemo-preventive properties of apigenin by suppressing angiogenesis and autophagy induction were also reported [13]. Further, apigenin repressed colon cancer migration and invasion by inhibiting the epithelial-mesenchymal transition/EMT) via NF-κB/Snail pathway [14]. However, all these are only the tip of the iceberg, and further effort is needed to describe the precise anti-cancerous mechanism of apigenin in CC.

In the current study, network pharmacology (a developing area of pharmacology for drug discovery) was utilised to explicate the possible mechanism of apigenin in CC by revealing apigenin’s relevant targets (in CC). Functional enrichment analysis further revealed the importance of the identified best targets. *In silico* methodologies such as molecular docking was utilised to investigate the interaction between apigenin and its targets in CC. Moreover, the identified targets (HSP90AA1, PTGS2, and MMP9) of apigenin were checked for their involvement in CC by studying their expression profile in CC and their impact on CC patient survival. Further, molecular dynamics simulation (MDS) analyses validated the protein-ligand interactions revealed by molecular docking. Finally, we checked the cytotoxicity of apigenin in human colon cancer cells (COLO-205). Influence of apigenin on the gene expression of the best target of apigenin (*HSP90AA1*) also investigated using colon cancer cells.

## 2. EXPERIMENTAL SECTION

### 2.1. Target Prediction

The targets of apigenin were divulged with the Swiss Target Prediction, STITCH database, and the PharmMapper database based on combined similarity measures (both 2-dimensional and 3-dimensional) with known ligands and pharmacophore model database [15-16]. The molecular structure of apigenin (in canonical SMILE format) was saved from PubChem [17] and used as input in different servers used in this study. The target protein names were changed to their corresponding gene symbol by utilising the database UniProt (https://www.uniprot.org/), and only “*Homo sapiens*” genes were selected.

In contrast, possible targets in CC were explored with the help of Online Mendelian Inheritance in Man (OMIM), GenCLiP3, GeneCards, and Therapeutic target database (TTD) databases by using keywords like “Colon Cancer”, “Colorectal Cancer”, and “CRC”. Shared targets of apigenin, and CC were acquired by plotting a Venn diagram employing an online tool (http://bioinformatics.psb.ugent.be/webtools/Venn/), further utilised in downstream experiments.

### 2.2. Protein-Protein Interaction (PPI) analysis and functional enrichment study

The protein interaction network of the targets of apigenin (in CC) was constructed with the help of the STRING database (v11.5) [18] and Cytoscape (v3.9.1) [19]. The shared/overlapping genes between the targets of apigenin and CC were used as input to develop the PPI network. In the case of PPI analyses, the species was restricted to only “*Homo sapiens*,” and the cut-off confidence score was set to > 0.70. The PPI network was further investigated for extremely interconnected regions (hub genes/core genes) by exploiting the MCODE application of Cytoscape (v3.9.1) by following parameters like a degree cut-off of 2, K-Core of 2, node score cut-off of 0.2, and a maximum depth of 100 [20]. The Gene Ontology (GO) analyses with the hub genes were executed through the ShinyGO server (v0.76) (http://bioinformatics.sdstate.edu/go/) to explore how these genes enriched in diverse Biological Processes, and Molecular Functions; KEGG and Reactome pathways. A term was considered significant for the experiments executed via ShinyGO if the FDR (false discovery rate) p-value < 0.05 [21].

### 2.3. Molecular docking analysis

The molecular interactions between the apigenin and its potential targets (in CC) were investigated by molecular docking with the AutoDock Vina software. RCSB PDB databank (https://www.rcsb.org/) was accessed to get the 3D structures of the relevant CC target proteins. On the other hand, the PubChem database was accessed to acquire the apigenin structure file (structure data file/SDF), converted to pdb format (by Open Babel 2.4.1) and finally saved in pdbqt format utilising AutoDock 4. The crystal structures of proteins in pdb format from the PDB database were saved. The preparation step of docking included cleaning up protein structure by deleting attached ligands, ions, and water molecules. Protein structures were further processed by attaching polar hydrogens, charges, assigning AD4 atom type with the help of AutoDock 4 to be eligible for docking. Finally, protein structures were saved in pdbqt format. AutoDock 4 was used to generate grid box coordinates which covered the whole protein structure. Discovery Studio Visualizer helped to inspect the protein-ligand interfaces [22].

### 2.4. Expression and genomic alteration data of apigenin targets in CC

The top target genes of apigenin in CC (revealed via docking score) were analysed to check differential gene expression in CC. The TNMplot database (https://www.tnmplot.com/), one of the most extensive cancer transcriptomic databases, was utilised to check differential gene expression for normal, tumour, and metastatic tissues [23]. The expression of the top targets in the protein level was also checked via the UALCAN server, which uses data from the CPTAC (Clinical Proteomic Tumor Analysis Consortium) dataset [24].

The genomic alterations of the top targets of apigenin were investigated within a Colon Cancer TCGA dataset (CPTAC-2 Prospective, Cell 2011), which includes 110 human colon adenocarcinoma tumour samples with paired normal samples. The CBioportal (https://www.cbioportal.org) server, which helps to study multidimensional cancer genomics data sets, was exercised to study genomic alterations [25].

### 2.5. Overall survival (OS) assessment

PrognoScan (http://dna00.bio.kyutech.ac.jp/PrognoScan/) database, which uses microarray data, was employed to reveal a possible correlation between gene expression (of hub genes) and patient survival in colon cancer (adjusted cox P value < 0.05) [26].

### 2.6. Molecular Dynamics Simulation (MDS) analyses

Besides the binding conformation and score from the molecular docking, the dynamic behaviour of the ligand binding with protein was studied using molecular dynamics simulation. The dynamic behaviour changes of apigenin with target proteins (selected after molecular docking analyses) were compared with control/reference inhibitors with the help of GROMACS 2019.6. The CHARMM27 force field converted PDB structures of protein-ligand complexes into GROMACS files [27]. Protein-ligand complexes were retained in a dodecahedron box by maintaining a distance (1.0 nm) from box edges and solvated by the TIP3P water model. The “genion” module of gromacs neutralised the complex system’s net charge with sodium and chlorine ions. The steepest descent algorithm accomplished system energy minimisation until the maximum force goes less than 10kJ/mol. Canonical ensemble and isobaric-isothermal ensemble were performed for 100 Pico-seconds with Velocity-rescale thermostat and Berendsen pressure coupling, respectively. Finally, the MDS run of the equilibrated protein-ligand complex system was executed with two Femto second (2 fs) time steps for 100 nanoseconds. Analyses of the stability and the strength of the protein-ligand complexes were performed in terms of MD parameters, including root means square deviation/RMSD, root means square fluctuations/RMSF, H-Bond numbers, and pair distance. Snapshots of the protein-ligand complexes were also extracted from the trajectory file to trace the dynamics of ligand molecules inside the binding pocket.

### 2.7. MMPBSA - binding free energy analysis

In addition to the stability analysis of the protein-ligand complex in a dynamic solvent environment, the strength of binding forces (regard to binding free energies) between protein and ligand was also evaluated using the MMPBSA method (Molecular Mechanics Poisson-Boltzmann Surface Area). MM-PBSA method calculated the binding free energy of complexes (protein-ligand) in terms of Van der Waals energy, Polar solvation and SASA energy, and Electrostatic energy from MD trajectories. The g_mmpbsa gromacs function was used to calculate the binding free energy of complex (protein-ligand) with a 1 ns time interval from MD trajectory [28]. The binding free energy (ΔGbind) of each complex was calculated using the equation described by Kollman et al., 2000 [29].

### 2.8. Cell culture and viability assay

Colon cancer cell line COLO-205 was obtained from the NCCS, Pune, cultured in RPMI with 10% foetal bovine serum and antibiotics (penicillin 50*μ*g/ml, streptomycin 50*μ*g/ml, and neomycin 100*μ*g/ml) within a humidified incubator (5% CO2 and temperature 37 °C). Apigenin (Sigma-Aldrich Co. purity of 99%) was dissolved in DMSO (dimethyl sulfoxide, HIMEDIA, India) (stored at −20 °C). Cell viability was measured using the MTT assay. 1 × 10^4 cells were treated with different concentration of apigenin (10-160 *μ*m) or control (DMSO) for 48 h and subsequently incubated with 20 μL of MTT (5mg/ml) at 37°C in the dark for 4 h [30]. A solubilisation solution (10% SDS in 0.01M HCL) was added to dissolve formazan crystals and further incubated at 37°C overnight. The absorbance at 570 nm was measured using a Thermo Scientific microplate reader.

### 2.9. Quantitative real-time PCR

The expression of the target genes was checked after total RNA extraction from cells treated with Apigenin or control DMSO using Trizol reagent (Invitrogen), followed by complementary DNAs (cDNAs) synthesis using a cDNA synthesis kit (BioBharati LifeScience, India) by following the manufacturer’s instructions. Expression was then quantified with the help of SYBR green PCR master mix (Thermo Scientific) and primer pair of *HSP90AA1* (F: 5’-AACTCAGACCCAGTCTTGTGGATGG-3’, R: 5’-AACCATCTCCTGCTAGCGCCTC-3’) and *ACTB* (F: 5’-GGATTCCTATGTGGGCGACGAG-3’; R: 5’-GAGCCACACGCAGCTCATTG-3’) with the help of an Applied Biosystem StepOne^™^ real-time PCR machine. Relative expression of *HSP90AA1* levels were determined following the 2^−ΔΔCT^ method based on C_T_ values, whereas *ACTB* gene expression was used for normalisation [31].

## 3. RESULTS AND DISCUSSION

### 3.1. Apigenin targets specific set of proteins related to colon carcinogenesis

172 unique targets of apigenin (after duplication removal) (Supplementary Table 1) and 442 CC-related target genes (Supplementary Table 2) were collected from different databases. The Venn diagram of these target gene sets were used to identify 49 shared/overlapped targets, representing different plausible targets of apigenin in CC (Supplementary Table 3). To explicit the relevance of these targets in CC, a PPI network was developed (via STRING), which was further processed with the Cytoscape, revealing a total of 47 nodes and 278 edges within the PPI network (Figure 1A). MCODE plugin of the Cytoscape application further identified highly interconnected regions (comprised of the hub genes) within the PPI network based on connectivity degree. The top-ranked MCODE cluster (score 10.833, k-core 2) comprised 13 nodes and 65 edges (Figure 1A) and was subjected to functional enrichment analyses. Gene ontology analyses with the ShinyGo server revealed that MCODE genes which represent apigenin’s core targets in CC are significantly related to different critical biological processes (e.g., “Response to UV”, “Response to chemical stress”, “Response to oxidative stress”, “Regulation of cell death”) (Figure 1B) and molecular functions (e.g., “Nitric-oxide synthase regulator activity”, “phosphatase binding”, “protein kinase binding” (Figure 1C). MCODE genes were also significantly enriched in different KEGG pathways, including “Colorectal cancer,” “Bladder cancer,” “MAPK signalling pathway,” and “MicroRNAs in cancer (Fig. 1D), linked to carcinogenesis. The Reactome pathway analysis further revealed that hub genes were enriched in several pathways linked to carcinogenesis, including ERBB2 signalling, signalling by non-receptor tyrosine kinases, PI3K/AKT signalling, and signalling by interleukins (Figure 1E). Thus, the top 13 targets of apigenin (e.g., VEGFA, HSP90AA1, TP53, HIF1A) were identified in CC, which plays an integral role in carcinogenesis, and could be targeted in CC.

**Table 1:** Summary of molecular docking analyses of apigenin and its target proteins in colon cancer. Reference compound of each target protein is represented in italics.

| Target protein with PDB ID | Ligand with PubChem CID | Binding energy (Kcal/mol) | No. of H-bond | H-bond forming residues | No. of Non-covalent interactions |
| --- | --- | --- | --- | --- | --- |
| HSP90AA1 (4BQG) | Apigenin (5280443) | -9.8 | 3 | D93, S52, L107 | 18 |
|  | *Geldanamycin* (5288382) | -6.6 | 2 | *R46, E47* | *10* |
| PTGS2 (5F19) | Apigenin (5280443) | -9.4 | 1 | M522 | 18 |
|  | *Ibuprofen* (3672) | -7.7 | 3 | *G526, L531* | *15* |
| MMP9 (1GKC) | Apigenin (5280443) | -9.2 | 2 | P421, R424 | 17 |
|  | *Mmp-9-IN-1* (135415473) | -9.2 | 3 | *A191, H401, H411* | *24* |

**Table 2:** Mean values of molecular dynamics simulation parameters of protein-ligand interactions over 100ns.

|  | HSP90 |  |  | MMP9 |  |  | PTGS2 |  |  |
| --- | --- | --- | --- | --- | --- | --- | --- | --- | --- |
|  | Apo protein | Apigenin | Control | Apo protein | Apigenin | Control | Apo protein | Apigenin | Control |
| **RMSD** | 0.1706 | 0.1237 | 0.1641 | 0.3701 | 0.2462 | 0.2981 | 0.2090 | 0.2096 | 0.1730 |
| **RMSF** | 0.1219 | 0.1161 | 0.1370 | 0.1849 | 0.1414 | 0.1385 | 0.1247 | 0.1229 | 0.1267 |
| **H-bond** | --- | 0.8297 | 0.6810 | --- | 0.2442 | 1.2440 | --- | 0.9530 | 2.083 |
| **Pair distance** | --- | 0.1793 | 0.2156 | --- | 0.1807 | 0.11902 | --- | 0.1885 | 0.1687 |

**Table 3:** Binding free energy analysis of apigenin and control inhibitors (Reference). with target proteins. References are indicated in italics.

|  | **HSP90** |  | **MMP9** |  | **PTGS2** |  |
| --- | --- | --- | --- | --- | --- | --- |
|  | **Apigenin** | *Geldanamycin* | **Apigenin** | *Mmp-9-IN-1* | **Apigenin** | *Ibuprofen* |
| **van der Waal energy (kJ/mol)** | -155.576 | -83.313 | -149.559 | -199.049 | -113.275 | -95.397 |
| **Electrostatic energy (kJ/mol)** | -19.242 | -38.880 | -0.558 | -58.479 | -16.345 | -194.184 |
| **Polar solvation energy (kJ/mol)** | 66.974 | 97.728 | 52.880 | 136.655 | 67.658 | 172.766 |
| **SASA energy (kJ/mol)** | -15.472 | -10.344 | -13.639 | -17.041 | -12.769 | -13.970 |
| **Binding energy (kJ/mol)** | -123.316 | -34.808 | -110.875 | -137.914 | -74.730 | -130.785 |

**Figure 1:**
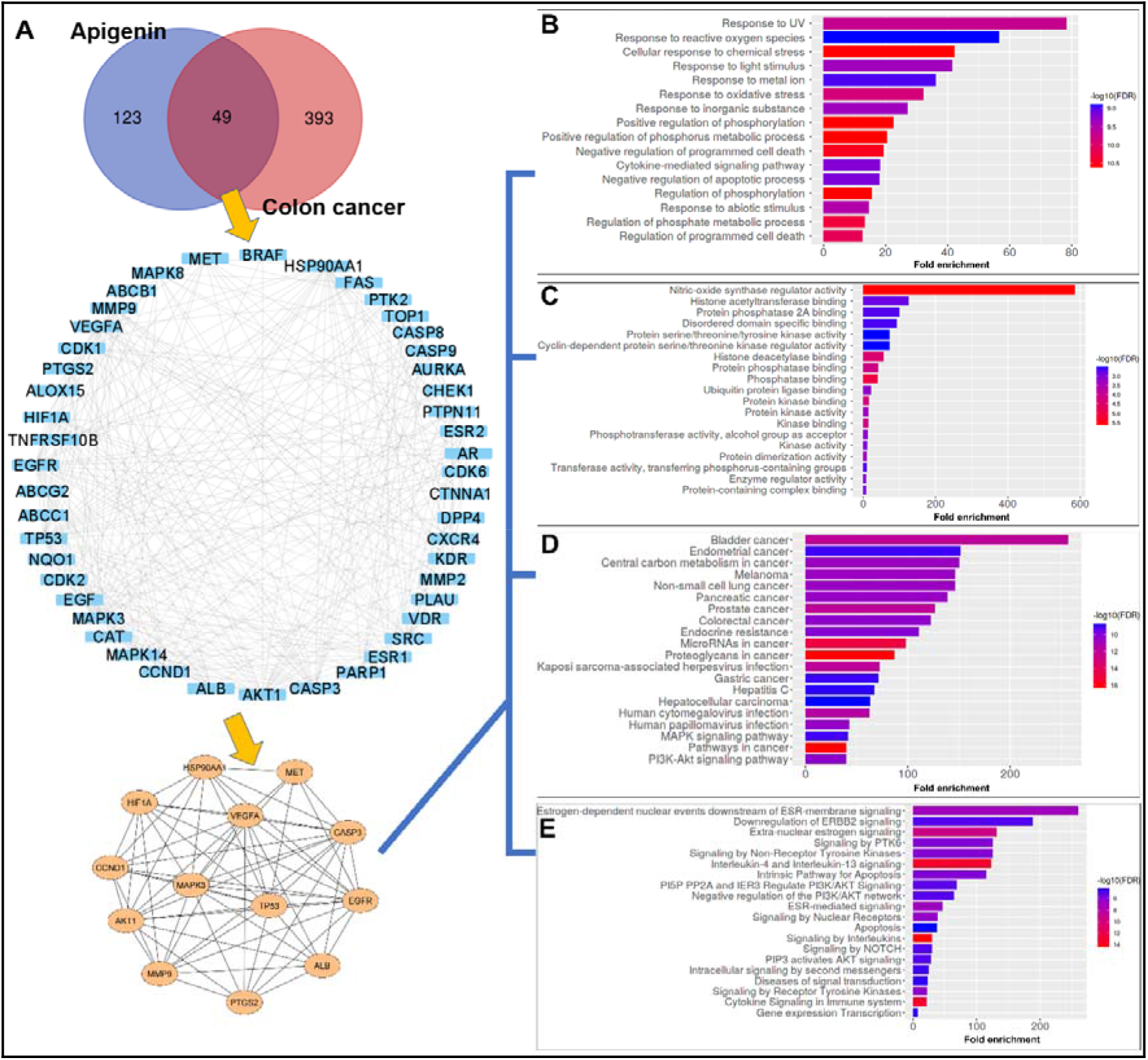
Protein-protein interaction (PPI) network of apigenin in colon cancer (CC). A) The PPI network of the overlapping targets between apigenin and CC and the highly interconnected region or core targets within the PPI. Functional enrichment analyses of the core targets of apigenin in CC, B) GO Biological processes, C) GO Molecular functions, D) KEGG pathways, and E) Reactome pathways.

### 3.3. Apigenin interacts strongly with its target proteins

The top 13 targets located in the core of the PPI network, as revealed by the MCODE, were subjected to molecular docking to validate apigenin’s targets in CC. The interaction analyses revealed that binding free energies of apigenin with target proteins ranged from -6 to -9.8 Kcal/mol. Heat shock protein 90 alpha family class A member 1 (HSP90AA1), prostaglandin-endoperoxidase synthase 2 (PTGS2), and matrix metalloproteinase 9 (MMP9) came out as the top 3 targets based on binding affinity, -9.8 Kcal/mol, -9.4 Kcal/mol, and -9.2 Kcal/mol respectively (Supplementary table 4). HSP90AA1 and apigenin interaction involved three H-bonds with residues including Ser52 (distance 3.16 Å), Asp93 (distance 4.61 Å), and Leu107 (distance 3.97 Å), along with several hydrophobic interactions (Fig. 2A). Hsp90 performs its chaperone activities by undergoing an ATPase cycle to aid in proper folding and maturation of client proteins [32]. Different amino acids form the enzyme’s active site (polar, charged, and hydrophobic amino acids), including Leu48, Asn51, Ser52, Asp93, Leu107, Phe138, Thr184 etc. The polar residue Asp93 is particularly important for Hsp90 action as it interacts directly with the ATP molecule [33]. In the docking interaction study, Asp93 was identified as one of the residues with which apigenin interacted via H-bonding (Fig. 2A), suggesting possible interference in Hsp90 action by apigenin. Apigenin also interacted with residues like Asn51, Ala55, Gly97, Asn106, and Phe138 (via non-bonded interactions), which lie on the crater towards the path of Asp93 (Fig. 2A). Thereby, apigenin could act as an ATP-competitive inhibitor of Hsp90 like geldanamycin (an established inhibitor of Hsp90). Interestingly, the binding free energy of the apigenin-HSP90AA1 interaction was much lower (-9.8 Kcal/mol) than the apigenin-reference compound (Geldanamycin) interaction (-6.6 Kcal/mol) which suggests a much stronger binding affinity of apigenin towards the HSP90AA1 (Table 1).

**Figure 2:**
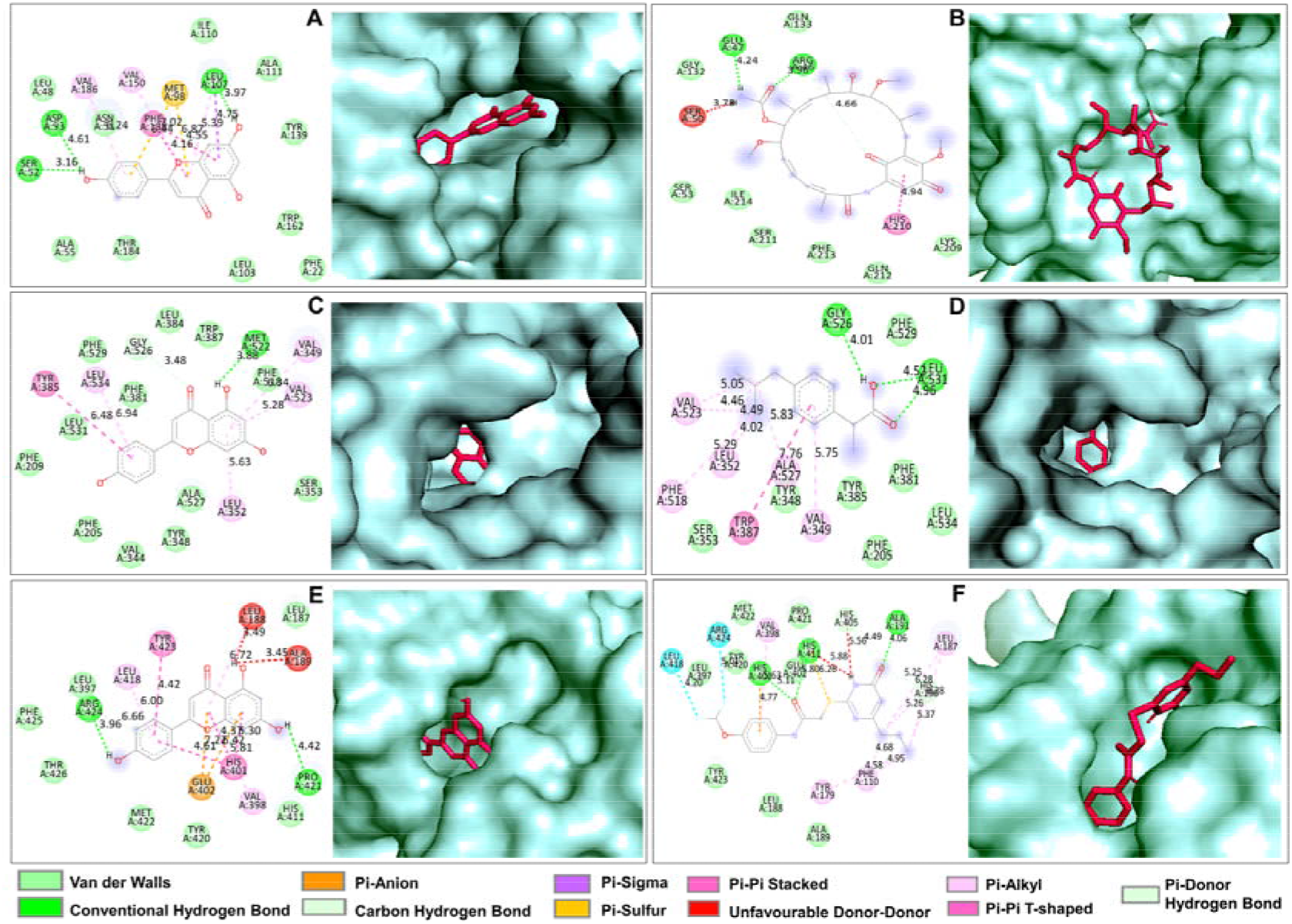
Molecular Docking interactions of apigenin-target proteins in colon cancer along with respective reference compound of target proteins. 2D and 3D interaction images of HSP90AA with A) apigenin and B) geldanamycin; PTGS2 with C) apigenin and D) ibuprofen; MMP9 with E) apigenin and F) Mmp-9-IN-1.

Apigenin also showed better binding affinity towards PTGS2 (COX-2) compared to the reference compound (Ibuprofen), as evidenced by the binding free energy of the protein-ligand interactions, -9.4 Kcal/mol and -7.7 Kcal/mol for apigenin-PTGS2 and ibuprofen-PTGS2 respectively (Table 1). COX enzymes (COX-1 and COX-2) regulate prostaglandins synthesis, which is crucial in the pathophysiology of inflammation [34]. Twenty-four amino acids line the active site of COX enzymes with only one amino acid difference between COX-1 and COX-2 (Ile523 to Val523) [35]. Apigenin was able to interact with several active site residues, including Phe205, Phe209, Val344, Tyr348, Val349, Leu352, Ser353, Phe381, Leu384, Tyr385, Trp387, Phe518, Val523, Gly526, Ala527, Leu531, and Leu534 by carbon-hydrogen bond, hydrophobic interactions, and Van der Walls interactions (Figure 2C). Among these interactions, Val523 and Phe518 are noteworthy as these two residues are part of the additional side pocket of COX-2 (absent in COX-1), which is crucial for COX-2 drug selectivity [36]. Thereby, apigenin could selectively block COX-2 action by interacting with the enzyme’s active site. On the other hand, apigenin interacted with the active site residue (Glu 402) and different metal binding sites (Leu 187, His401, His411) of the MMP-9 enzyme with an excellent binding affinity (-9.2 Kcal/mol) (Table 1). The apigenin-MMP-9 interaction was stabilised by hydrogen bonds and different non-bonded interactions (Figure 2E). The binding affinity of apigenin towards the MMP-9 was similar to the Mmp-9-IN-1 (reference compound) based on the binding free energy (Table 1), suggesting possible inhibition of MMP-9 enzymatic action by apigenin.

### 3.4. Elevated expression of top targets is observed in colon cancer

Altered gene expression is pivotal at the beginning of cancer and throughout the cancer progression. Therefore, we checked gene expressions of the top targets of apigenin in colon cancer. The gene expression level of the top targets using the TNMplot database showed that *HSP90AA1* expression increased significantly by 1.29-fold in colon adenocarcinoma, COAD (Figure 3A). Similarly, *PTGS2* and *MMP9* expression also increased in colon tumour tissues compared to the normal tissues (median fold change 2.06 and 2.09, respectively) (Figure 3B and 3C). Protein expression data from the UALCAN database also showed that all three targets were overexpressed in tumour tissues compared to normal tissues in colon cancer (Figure 3D, 3E, and 3F). Therefore, these three proteins might have a crucial role in colon cancer and targeting these proteins with apigenin would be crucial in search CC therapeutics. The top targets of apigenin were found to be altered in genomic level of 22% of patients (N = 110) within a TCGA dataset (colon cancer, CPTAC-2 Prospective, Cell 2019). *MMP9* alteration frequency was highest (11%) in CC among the three genes, followed by *HSP90AA1* (8%) and *PTGS2* (5%), respectively, which includes mutations and copy number alterations (Figure 3G, 3H and 3I). Higher expression of the *HSP90AA1, MMP9*, and *PTGS2* was also correlated to poor overall survival (OS) in CC, based on adjusted cox P-value, revealed from the PrognoScan server. Expression of the top targets was positively associated with CRC overall survival in the case of *HSP90AA1* (Figure 5A; VMC cohort; corrected *P*-value = 0.004) (Figure 3J). On the other hand, high expression of *PTGS2* and *MMP9* also tends to be correlated with poor OS (Figure 3K and 3L), however, these two observations were not statistically significant. Overall, expression analyses, genomic alterations and survival studies of these three targets suggest their contribution to colon tumorigenesis.

**Figure 3:**
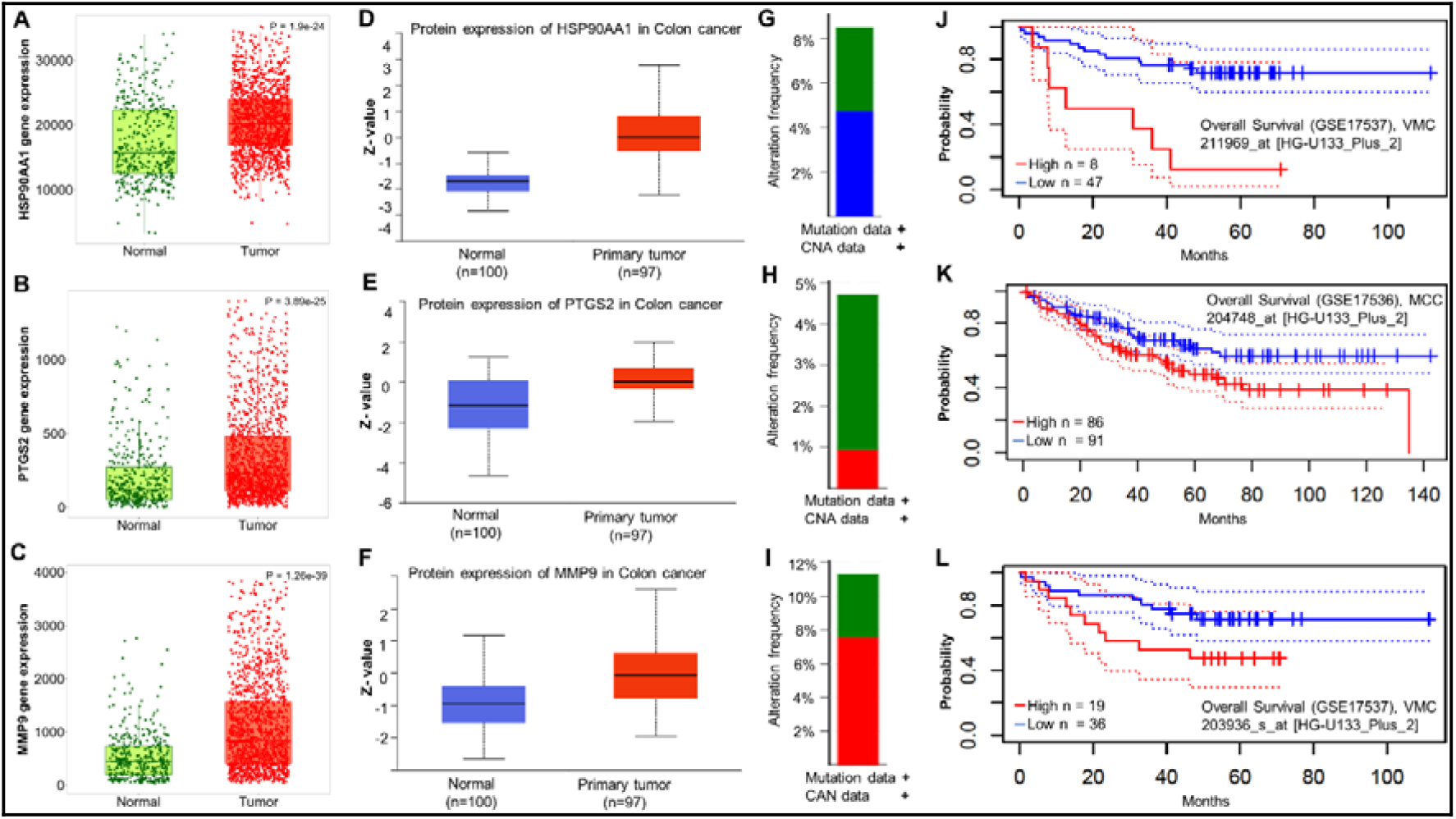
Higher genetic alteration and gene expression of top thee targets of Apigenin correlate with lower survival of the CC patients. Expression of the top 3 targets of apigenin in colon cancer, mRNA, and protein expression of HSP90AA1 (A & D), PTGS2 (B & E), and MMP9 (C & F). Genomic alteration frequency of the top target genes of apigenin in colon cancer from CBioportal (CPTAC-2 dataset), *HSP90AA1* (G), *PTGS2* (H), and *MMP9* (I). The survival curve (Kaplan-Meier) for high (red) and low (blue) gene expression groups from different datasets was used to analyse the prognostic value of *HSP90AA1* (J), *PTGS2* (K), and *MMP9* (L) in colorectal cancer (PrognoScan server).

### 3.5. Characterization of Protein-ligand interaction by MD simulation

The dynamic behaviour of apigenin with target proteins was analysed in a solvated environment with the help of molecular dynamics simulation to achieve in depth understanding of the stability and strength of the interactions. MDS provides the platform to mimic cellular or *in vitro* experimental conditions through which the binding of ligands with target proteins can be analysed efficiently. This study analysed the strength and the stability of apigenin, and target proteins based on RMSD (root means square deviation), RMSF (root means square fluctuations), number of H-bonds and pair distance. The average values of the MD parameters were tabulated in Table 2.

The root means square deviations of the protein backbones give the detail about the protein stability upon ligand binding. More deviation in the protein backbone indicates a less stable structure; however, the binding of apigenin caused less deviation in HSP90 and MMP9 structure compared to the control inhibitor and unbound state. In the case of PTGS2, both unbound and apigenin complexes deviated at the same level between 0.1 to 0.2 nm range, and the control inhibitor showed lesser deviation (Figure 4A). The residual level fluctuation over 100 ns of target proteins with and without inhibitors shed light on residues involved in the interactions. Residues from 60 to 75 of both HSP90 and MMP9 fluctuated differently compared to the unbound state. These are nearby residues of the binding pocket, which fluctuated more by the ligand binding to accommodate them in the binding pocket (Figure 4B). In the case of PTGS2, the residual fluctuations are negligible, as is the difference between the average RMSFs (Table 2). To conclude, the binding of apigenin with target proteins did not destabilise the protein backbone. Besides the protein stability, the strength of the interactions was analysed based on the number of H-bonds formed and the distance of the pairing between inhibitor and protein. Among three target proteins, apigenin formed 0 to 5 H-bonds with HSP90 and PTGS2 and 0 to 2 H-bonds with MMP9 (Figure 5). In addition to that, the pair distance between apigenin and target proteins is also lesser than 0.18 nm over 100 ns simulation (Table 2), which is good enough to hold the ligand molecules inside the binding pocket. However, the control inhibitors of MMP9 and PTGS2 exhibited better activity than apigenin in the average H-bonds formed and pair distance over 100 ns (Table 2). Based on the binding strength analysis, apigenin can establish stronger binding than control with HSP90 and comparatively lesser than control inhibitors with MMP9 and PTGS2. The contact analysis of the inhibitors was also performed to trace the movement of inhibitors inside the binding pocket by extracting the structures from the MD trajectory. It has been observed that apigenin and control inhibitors of all target proteins remained in the same binding pocket throughout the simulation with minor conformation changes except the control inhibitor of HSP90 (Figure 6). The change in the binding pocket was observed for the control inhibitor of HSP90 after 40th ns, which can also be observed in the pair distance graph of the same (Figure 6).

**Figure 4:**
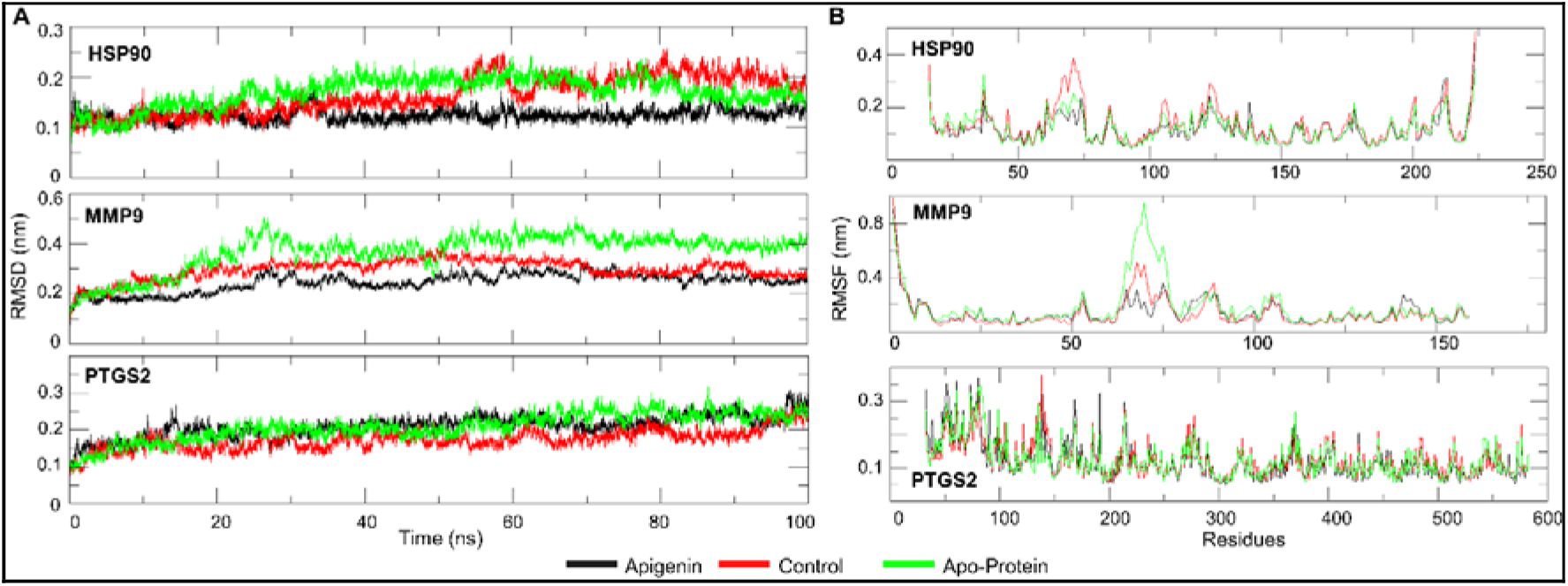
Root-mean-square deviation (RMSD) and Root means square fluctuation (RMSF) of the backbone of protein-ligand complexes were plotted over 100 ns Molecular Dynamics Simulation. A) RMSD of protein backbone upon binding of apigenin and control inhibitors over 100 ns. B) RMSF of each residue of target proteins upon binding of apigenin and control inhibitors over 100 ns.

**Figure 5:**
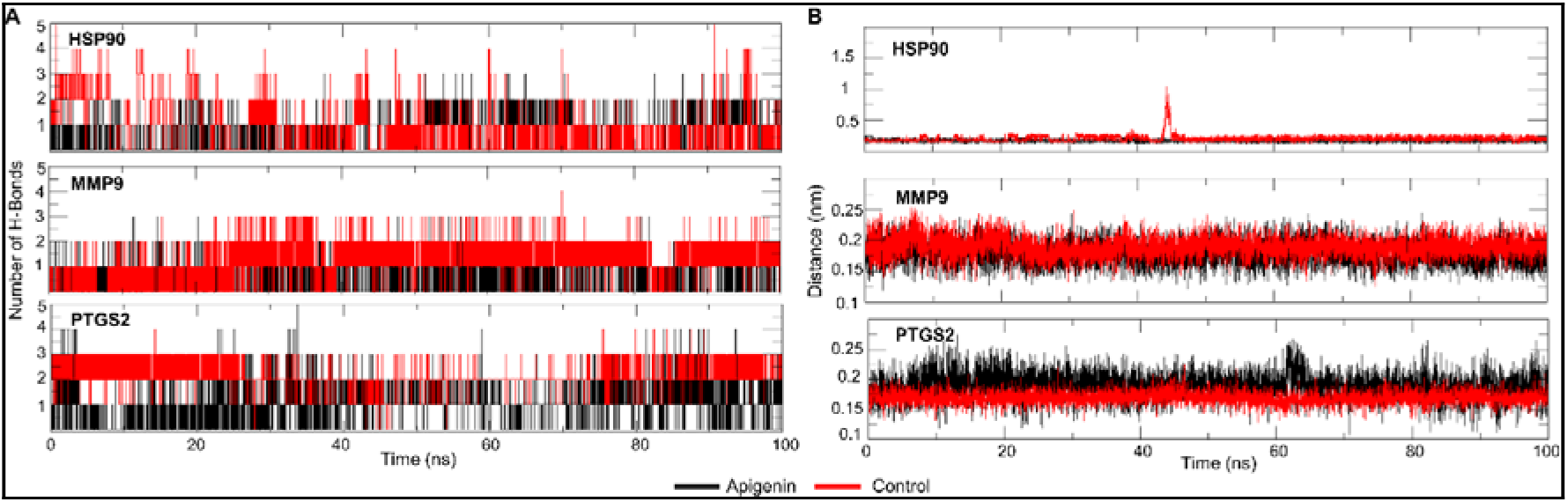
Inter-molecular hydrogen bonds, and Hydrogen bond pair distance between target proteins and candidate drugs over 100 ns MD simulation. A) Number of Hydrogen bonds formed by the apigenin and control inhibitors with target proteins over 100 ns time-period. B) Distance maintained by the apigenin and control inhibitors with target proteins upon binding over 100 ns simulation.

**Figure 6:**
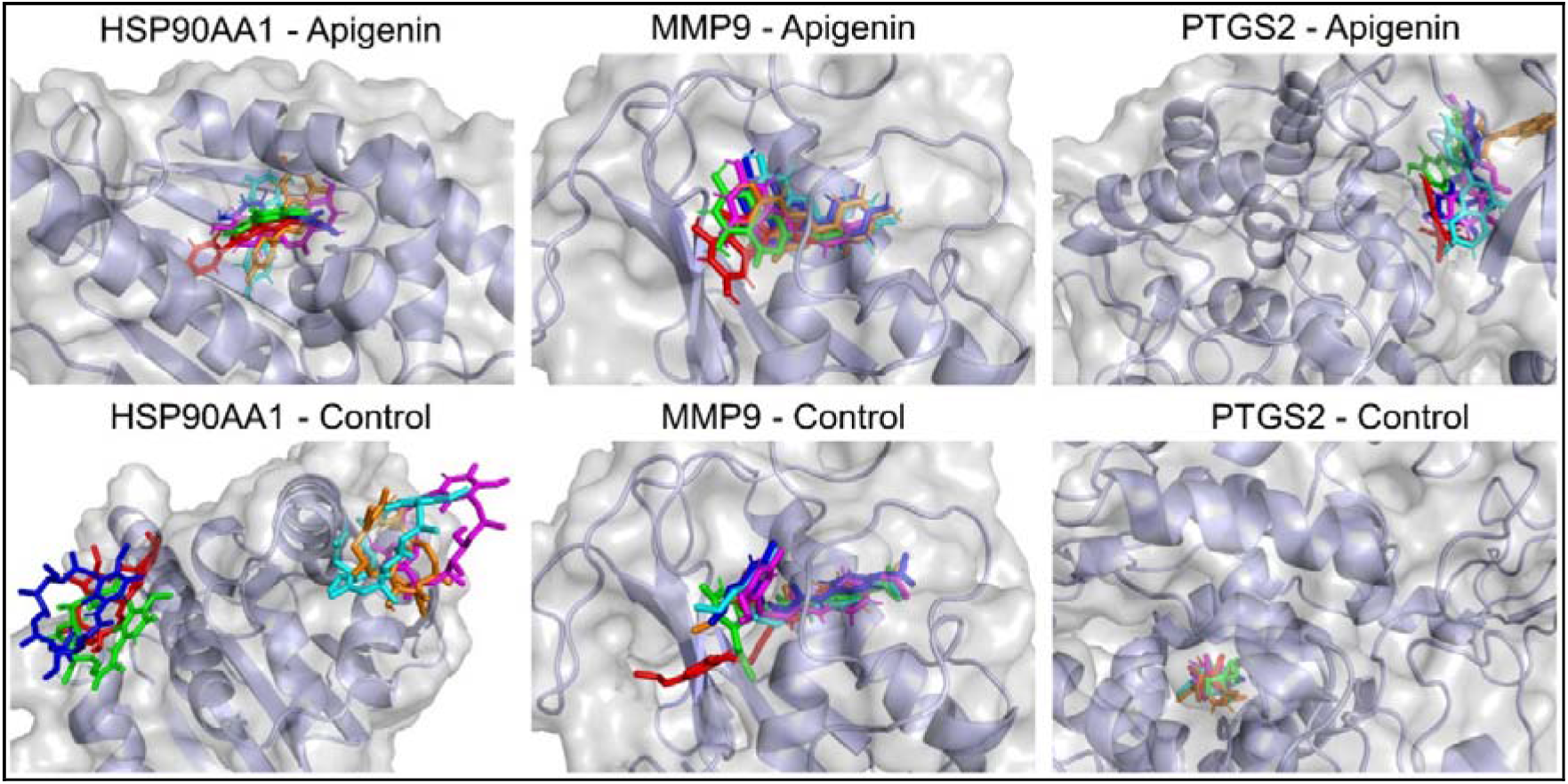
Dynamics of the top three target protein of apigenin in colon cancer on the surface of target proteins during Molecular Dynamics Simulation (100 ns). Binding conformation of apigenin and control inhibitors in the binding pockets of target proteins with 20 ns time interval. The ligand molecules are depicted in different colours inside the binding pocket based on the time frame, red- 0^th^ ns, green-20^th^ ns, blue- 40^th^ ns, magenta- 60^th^ ns, cyan- 80^th^ ns, orange- 100^th^ ns.

### 3.6. Binding Free energy analysis selected the best target of apigenin in colon cancer

Intermolecular forces involved in the protein-ligand interaction are essential in defining the binding strength. In this study, the forces involved in ligand interactions, such as Van der Waal, electrostatic, polar solvation and SASA, were calculated using MMPBSA binding free energy method. The binding free energies were calculated from the MD trajectories with one ns time interval and tabulated in Table 3. Apigenin attained the lowest binding free energy with HSP90AA1 (-123.316 kJ/mol), which is better than the HSP90AA1 control/reference compound also, followed by MMP9 (-110.875 kJ/mol) and PTGS2 (-74.730 kJ/mol). In the case of MMP9 and PTGS2, control inhibitors exhibited better binding energy than apigenin due to the lowest electrostatic energy of apigenin with target proteins. The difference between the binding energy of apigenin and control with MMP9 is also comparable. Following MDS and binding free energy analysis, apigenin can be considered as a potent inhibitor for HSP90AA1, followed by MMP9 and PTGS2, respectively.

### 3.7. Apigenin reduced COLO-205, colon cancer cell viability and reduced the mRNA expression of HSP90AA1

Apigenin’s anti-proliferative or cytotoxic effects were investigated in human colon cancer cells (COLO-205) by treating cells with different concentrations of Apigenin (10-160 µm) for 48 h (Figure 7). The viability of COLO-205 cells was significantly reduced in a dose-dependent manner in the presence of Apigenin. As revealed by the MTT assay, Apigenin at lower concentrations (10-40*μ*M) reduced cell viability by 10-20% in comparison to untreated group. In contrast, higher concentrations of Apigenin (60-160*μ*M) reduced cell viability by almost 50% (Figure 7A). Next, we wanted to check whether Apigenin modulates the mRNA expression of HSP90AA1, as colon cancer cells depend on elevated Hsp90 for survival. Utilising Real-Time PCR, mRNA expression was checked in COLO-205 cells, and relative expression of the *HSP90AA1* was significantly reduced by up to 60% compared to untreated cells in a dose-dependent manner up to 40 *μ*M of apigenin exposure (Figure 7B). Additionally, in the case of higher concentrations of apigenin exposure, mRNA expression was significantly reduced to 50% (60*μ*M and 80*μ*M) or 40% (120*μ*M). Interestingly, a little upward shift in *HSP90AA1* expression was observed in the case of higher doses of Apigenin (60-120*μ*M) compared to lower doses (20*μ*M-40*μ*M) (Figure 7B). However, these differences in observations (between low and high apigenin groups) were not statistically significant.

**Figure 7:**
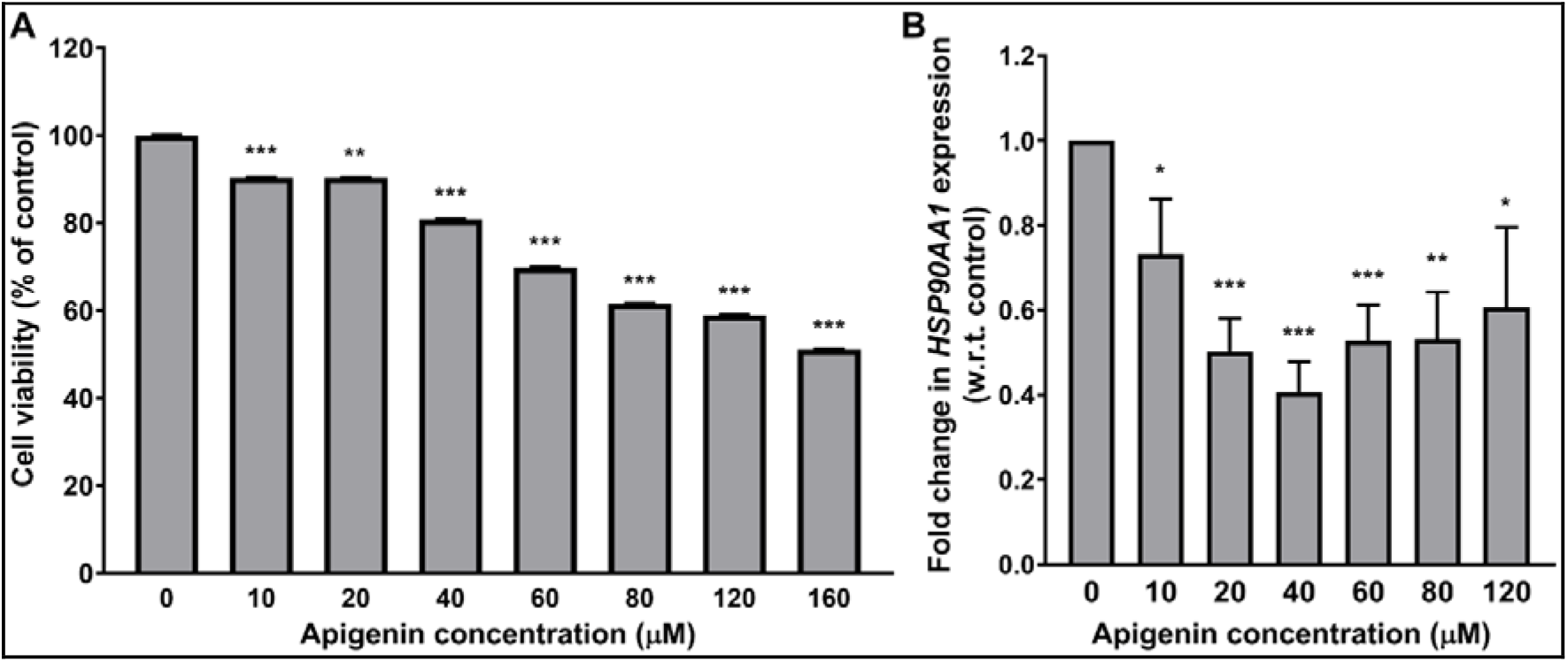
Effect of apigenin on COLO-205 cell viability and HSP90AA1 mRNA expression. **A)** Cell viability was reduced significantly upon apigenin exposure, and B) expression of HSP90AA1 mRNA diminished by apigenin. (\**P* < 0.05, \*\**P* < 0.01, \*\*\**P* < 0.001 vs untreated group, data represented as the mean ± SEM of at least three independent experiments).

Hsp90 protects against augmented cellular stress and creates a tumour-promoting environment during neoplastic transformation. In transformed cells, Hsp90 performs a unique role in preserving malignant phenotype by facilitating the accumulation of beneficial mutations and suppressing mutations which could be lethal for cancer cells [37]. HSP90AA1 expression is augmented in colon cancer as revealed by gene expression analysis (Figure 3), providing survival benefits to cancer cells. In contrast, Apigenin markedly inhibited *HSP90AA1* expression to reduce cancer cell (COLO-205) viability, which might be further cashed upon to develop a therapeutic intervention in colon cancer.

## 3. Conclusion

Apigenin, a natural flavone comes into the spotlight in recent times as an anti-cancer agent. However, the precise mechanism of apigenin action in cancer, particularly colon cancer, is still elusive. This study explored apigenin’s potential targets/pathways in colon cancer. With the help of available online resources and software, including STRING, Cytoscape, molecular docking, and molecular dynamics simulation, HSP90AA1, MMP9, and PTGS2 were revealed as top hits of apigenin in CC. Moreover, the best target of apigenin in CC, HSP90AA1, was checked for mRNA expression in colon cancer cells (COLO-205), and apigenin diminished the mRNA expression of HSP90AA1 and successfully reduced cancer cell viability. To the best of our knowledge, this study first reports HSP90AA1 as a target of apigenin in CC. These findings could be the stepping stone to understanding apigenin’s interactions in CC and developing novel therapeutic approaches.

## References

1. Petre-Mandache, C.B.; Margaritescu, D. N. Risk Factors and Genetic Predisposition in Colorectal Cancer: A Study on Young and Old Adults. Current Health Sciences Journal 2021, No. 1, 84–88. https://doi.org/10.12865/CHSJ.47.01.13.

2. Rawla, P.; Sunkara, T.; Barsouk, A. Epidemiology of Colorectal Cancer: Incidence, Mortality, Survival, and Risk Factors. pg 2019, 14 (2), 89–103. https://doi.org/10.5114/pg.2018.81072.

3. Siegel, R. L.; Miller, K. D.; Jemal, A. Colorectal Cancer Mortality Rates in Adults Aged 20 to 54 Years in the United States, 1970-2014. JAMA 2017, 318 (6), 572. https://doi.org/10.1001/jama.2017.7630.

4. Bhandari, A.; Woodhouse, M.; Gupta, S. Colorectal Cancer Is a Leading Cause of Cancer Incidence and Mortality among Adults Younger than 50 Years in the USA: A SEER-Based Analysis with Comparison to Other Young-Onset Cancers. J Investig Med 2017, 65 (2), 311–315. https://doi.org/10.1136/jim-2016-000229.

5. Shankar, E.; Kanwal, R.; Candamo, M.; Gupta, S. Dietary Phytochemicals as Epigenetic Modifiers in Cancer: Promise and Challenges. Seminars in Cancer Biology 2016, 40–41, 82–99. https://doi.org/10.1016/j.semcancer.2016.04.002.

6. Oruç, Z.; Kaplan, M. A. Effect of Exercise on Colorectal Cancer Prevention and Treatment. WJGO 2019, 11 (5), 348–366. https://doi.org/10.4251/wjgo.v11.i5.348.

7. Xie, Y.-H.; Chen, Y.-X.; Fang, J.-Y. Comprehensive Review of Targeted Therapy for Colorectal Cancer. Sig Transduct Target Ther 2020, 5 (1), 22. https://doi.org/10.1038/s41392-020-0116-z.

8. Zhao, Y.; Hu, X.; Zuo, X.; Wang, M. Chemopreventive Effects of Some Popular Phytochemicals on Human Colon Cancer: A Review. Food Funct. 2018, 9 (9), 4548–4568. https://doi.org/10.1039/C8FO00850G.

9. Ghi[u, A.; Schwiebs, A.; Radeke, H. H.; Avram, S.; Zupko, I.; Bor, A.; Pavel, I. Z.; Dehelean, C. A.; Oprean, C.; Bojin, F.; Farcas, C.; Soica, C.; Duicu, O.; Danciu, C. A Comprehensive Assessment of Apigenin as an Antiproliferative, Proapoptotic, Antiangiogenic and Immunomodulatory Phytocompound. Nutrients 2019, 11 (4), 858. https://doi.org/10.3390/nu11040858.

10. Imran, M.; Aslam Gondal, T.; Atif, M.; Shahbaz, M.; Batool Qaisarani, T.; Hanif Mughal, M.; Salehi, B.; Martorell, M.; Sharifi[Rad, J. Apigenin as an Anticancer Agent. Phytotherapy Research 2020, 34 (8), 1812–1828. https://doi.org/10.1002/ptr.6647.

11. Javed, Z.; Sadia, H.; Iqbal, M. J.; Shamas, S.; Malik, K.; Ahmed, R.; Raza, S.; Butnariu, M.; Cruz-Martins, N.; Sharifi-Rad, J. Apigenin Role as Cell-Signaling Pathways Modulator: Implications in Cancer Prevention and Treatment. Cancer Cell Int 2021, 21 (1), 189. https://doi.org/10.1186/s12935-021-01888-x.

12. Xu, M.; Wang, S.; Song, Y.; Yao, J.; Huang, K.; Zhu, X. Apigenin Suppresses Colorectal Cancer Cell Proliferation, Migration and Invasion via Inhibition of the Wnt/β-Catenin Signaling Pathway. Oncology Letters 2016, 11 (5), 3075–3080. https://doi.org/10.3892/ol.2016.4331.

13. Sung, B.; Chung, H. Y.; Kim, N. D. Role of Apigenin in Cancer Prevention via the Induction of Apoptosis and Autophagy. J Cancer Prev 2016, 21 (4), 216–226. https://doi.org/10.15430/JCP.2016.21.4.216.

14. Tong, J.; Shen, Y.; Zhang, Z.; Hu, Y.; Zhang, X.; Han, L. Apigenin Inhibits Epithelial-Mesenchymal Transition of Human Colon Cancer Cells through NF-?B/Snail Signaling Pathway. Bioscience Reports 2019, 39 (5), BSR20190452. https://doi.org/10.1042/BSR20190452.

15. Daina, A.; Michielin, O.; Zoete, V. SwissTargetPrediction: Updated Data and New Features for Efficient Prediction of Protein Targets of Small Molecules. Nucleic Acids Research 2019, 47 (W1), W357–W364. https://doi.org/10.1093/nar/gkz382.

16. Wang, X.; Shen, Y.; Wang, S.; Li, S.; Zhang, W.; Liu, X.; Lai, L.; Pei, J.; Li, H. PharmMapper 2017 Update: A Web Server for Potential Drug Target Identification with a Comprehensive Target Pharmacophore Database. Nucleic Acids Research 2017, 45 (W1), W356–W360. https://doi.org/10.1093/nar/gkx374.

17. Kim, S.; Chen, J.; Cheng, T.; Gindulyte, A.; He, J.; He, S.; Li, Q.; Shoemaker, B. A.; Thiessen, P. A.; Yu, B.; Zaslavsky, L.; Zhang, J.; Bolton, E. E. PubChem in 2021: New Data Content and Improved Web Interfaces. Nucleic Acids Research 2021, 49 (D1), D1388–D1395. https://doi.org/10.1093/nar/gkaa971.

18. Szklarczyk, D.; Gable, A. L.; Nastou, K. C.; Lyon, D.; Kirsch, R.; Pyysalo, S.; Doncheva, N. T.; Legeay, M.; Fang, T.; Bork, P.; Jensen, L. J.; von Mering, C. The STRING Database in 2021: Customizable Protein–Protein Networks, and Functional Characterization of User-Uploaded Gene/Measurement Sets. Nucleic Acids Research 2021, 49 (D1), D605–D612. https://doi.org/10.1093/nar/gkaa1074.

19. Shannon, P.; Markiel, A.; Ozier, O.; Baliga, N. S.; Wang, J. T.; Ramage, D.; Amin, N.; Schwikowski, B.; Ideker, T. Cytoscape: A Software Environment for Integrated Models of Biomolecular Interaction Networks. Genome Res. 2003, 13 (11), 2498–2504. https://doi.org/10.1101/gr.1239303.

20. Ren, C.; Li, M.; Du, W.; Lü, J.; Zheng, Y.; Xu, H.; Quan, R. Comprehensive Bioinformatics Analysis Reveals Hub Genes and Inflammation State of Rheumatoid Arthritis. BioMed Research International 2020, 2020, 1–13. https://doi.org/10.1155/2020/6943103.

21. Ge, S. X.; Jung, D.; Yao, R. ShinyGO: A Graphical Gene-Set Enrichment Tool for Animals and Plants. Bioinformatics 2020, 36 (8), 2628–2629. https://doi.org/10.1093/bioinformatics/btz931.

22. Trott, O.; Olson, A. J. AutoDock Vina: Improving the Speed and Accuracy of Docking with a New Scoring Function, Efficient Optimization, and Multithreading. J. Comput. Chem. 2009, NA-NA. https://doi.org/10.1002/jcc.21334.

23. Bartha, Á.; Győrffy, B. TNMplot.Com: A Web Tool for the Comparison of Gene Expression in Normal, Tumor and Metastatic Tissues. IJMS 2021, 22 (5), 2622. https://doi.org/10.3390/ijms22052622.

24. Chen, F.; Chandrashekar, D. S.; Varambally, S.; Creighton, C. J. Pan-Cancer Molecular Subtypes Revealed by Mass-Spectrometry-Based Proteomic Characterization of More than 500 Human Cancers. Nat Commun 2019, 10 (1), 5679. https://doi.org/10.1038/s41467-019-13528-0.

25. Gao, J.; Aksoy, B. A.; Dogrusoz, U.; Dresdner, G.; Gross, B.; Sumer, S. O.; Sun, Y.; Jacobsen, A.; Sinha, R.; Larsson, E.; Cerami, E.; Sander, C.; Schultz, N. Integrative Analysis of Complex Cancer Genomics and Clinical Profiles Using the CBioPortal. Sci. Signal. 2013, 6 (269). https://doi.org/10.1126/scisignal.2004088.

26. Mizuno, H.; Kitada, K.; Nakai, K.; Sarai, A. PrognoScan: A New Database for Meta-Analysis of the Prognostic Value of Genes. BMC Med Genomics 2009, 2 (1), 18. https://doi.org/10.1186/1755-8794-2-18.

27. Hart, K.; Foloppe, N.; Baker, C. M.; Denning, E. J.; Nilsson, L.; MacKerell, A. D. Optimization of the CHARMM Additive Force Field for DNA: Improved Treatment of the BI/BII Conformational Equilibrium. J. Chem. Theory Comput. 2012, 8 (1), 348–362. https://doi.org/10.1021/ct200723y.

28. Kumari, R.; Kumar, R.; Open Source Drug Discovery Consortium; Lynn, A. G_mmpbsa--a GROMACS Tool for High-Throughput MM-PBSA Calculations. J Chem Inf Model 2014, 54 (7), 1951–1962. https://doi.org/10.1021/ci500020m.

29. Kollman, P. A.; Massova, I.; Reyes, C.; Kuhn, B.; Huo, S.; Chong, L.; Lee, M.; Lee, T.; Duan, Y.; Wang, W.; Donini, O.; Cieplak, P.; Srinivasan, J.; Case, D. A.; Cheatham, T. E. Calculating Structures and Free Energies of Complex Molecules: Combining Molecular Mechanics and Continuum Models. Acc Chem Res 2000, 33 (12), 889–897. https://doi.org/10.1021/ar000033j.

30. van Tonder, A.; Joubert, A. M.; Cromarty, A. D. Limitations of the 3-(4,5-Dimethylthiazol-2-Yl)-2,5-Diphenyl-2H-Tetrazolium Bromide (MTT) Assay When Compared to Three Commonly Used Cell Enumeration Assays. BMC Res Notes 2015, 8 (1), 47. https://doi.org/10.1186/s13104-015-1000-8.

31. Livak, K. J.; Schmittgen, T. D. Analysis of Relative Gene Expression Data Using534119v1 Real-Time Quantitative PCR and the 2−ΔΔCT Method. Methods 2001, 25 (4), 402–408. https://doi.org/10.1006/meth.2001.1262.

32. Pearl, L. H. Review: The HSP90 Molecular Chaperone—an Enigmatic ATPase. Biopolymers 2016, 105 (8), 594–607. https://doi.org/10.1002/bip.22835.

33. Abbasi, M.; Sadeghi-Aliabadi, H.; Amanlou, M. Prediction of New Hsp90 Inhibitors Based on 3,4-Isoxazolediamide Scaffold Using QSAR Study, Molecular Docking and Molecular Dynamic Simulation. DARU J Pharm Sci 2017, 25 (1), 17. https://doi.org/10.1186/s40199-017-0182-0.

34. Choi, S.-H.; Aid, S.; Bosetti, F. The Distinct Roles of Cyclooxygenase-1 and -2 in Neuroinflammation: Implications for Translational Research. Trends in Pharmacological Sciences 2009, 30 (4), 174–181. https://doi.org/10.1016/j.tips.2009.01.002.

35. Zarghi, A.; Arfaei, S. Selective COX-2 Inhibitors: A Review of Their Structure-Activity Relationships. Iran J Pharm Res 2011, 10 (4), 655–683.

36. Oniga, S. D.; Pacureanu, L.; Stoica, C. I.; Palage, M. D.; Crăciun, A.; Rusu, L. R.; Crisan, E.-L.; Araniciu, C. COX Inhibition Profile and Molecular Docking Studies of Some 2-(Trimethoxyphenyl)-Thiazoles. Molecules 2017, 22 (9), 1507. https://doi.org/10.3390/molecules22091507.

37. Szczuka, I.; Wierzbicki, J.; Serek, P.; Szczęśniak-Sięga, B. M.; Krzystek-Korpacka, M. Heat Shock Proteins HSPA1 and HSP90AA1 Are Upregulated in Colorectal Polyps and Can Be Targeted in Cancer Cells by Anti-Inflammatory Oxicams with Arylpiperazine Pharmacophore and Benzoyl Moiety Substitutions at Thiazine Ring. Biomolecules 2021, 11 (11), 1588. https://doi.org/10.3390/biom11111588.

